# Differential CheR affinity for chemoreceptor C-terminal pentapeptides biases chemotactic responses

**DOI:** 10.1101/2024.03.07.583918

**Authors:** Félix Velando, Elizabet Monteagudo-Cascales, Miguel A. Matilla, Tino Krell

## Abstract

The capacity of chemotaxis pathways to respond to signal gradients relies on adaptation mediated by the coordinated action of CheR methyltransferases and CheB methylesterases. Many chemoreceptors contain a C-terminal pentapeptide at the end of a linker. In *Escherichia coli,* this pentapeptide forms a high-affinity binding site for CheR and phosphorylated CheB, and its removal interferes with adaptation. The analysis of all available chemoreceptor sequences showed that pentapeptide sequences vary greatly, and bacteria often possess multiple chemoreceptors that differ in their pentapeptide sequences. Using the phytopathogen *Pectobacterium atrosepticum* SCRI1043, we assessed whether this sequence variation alters CheR affinity and chemotaxis. SCRI1043 has 36 chemoreceptors, of which 19 possess a C-terminal pentapeptide. Using isothermal titration calorimetry, we show that the affinity of CheR for the different pentapeptides varies up to 11-fold (*K*_D_ of 90 nM to 1 µM). The pentapeptides with the highest and lowest affinities differed only in a single amino acid. Deletion of the *cheR* gene abolishes chemotaxis. PacC is the sole chemoreceptor for L-Asp in SCRI1043, and the replacement of its pentapeptide with those having the highest and lowest affinities significantly interfered with L-Asp chemotaxis. Variable pentapeptide sequences thus provide a mechanism to bias the responses mediated by chemoreceptors.

## Introduction

Bacterial chemotaxis is the directed swimming of bacteria in chemical gradients. It facilitates the migration of bacteria to sites that are favorable for survival. A major benefit from chemotaxis is access to nutrients (Colin *et al*., 2021; Matilla *et al*., 2023). For many human and plant pathogens with different lifestyles and infection mechanisms, chemotaxis is essential for virulence (Matilla and Krell, 2018). Chemotaxis signaling pathways are among the most abundant prokaryotic signal transduction mechanisms (Wuichet and Zhulin, 2010; Gumerov *et al*., 2023).

Chemotactic responses are typically initiated by a molecule binding to an extracytosolic sensor domain of a chemoreceptor. This binding generates a conformational change that is transmitted across the membrane to modulate the activity of the autokinase CheA, which in turn phosphorylates the CheY response regulator. Only the phosphorylated form of CheY is able to bind to the flagellar motor to control its direction and/or speed of rotation, ultimately resulting in chemotaxis (Parkinson *et al*., 2015; Bi and Sourjik, 2018).

The capacity of chemotaxis pathways to respond to signal gradients rather than to a constant signal concentration relies on adaptation mechanisms. Canonical adaptation is based on the coordinated action of the CheR methyltransferase and the CheB methylesterase, which catalyze the methylation and demethylation, respectively, of specific glutamyl residues in the chemoreceptor signaling domain (Parkinson *et al*., 2015; Bi and Sourjik, 2018). The importance of this adaptation mechanism is illustrated by the fact that CheR and CheB are highly conserved proteins that are present in almost all chemosensory pathways (Wuichet and Zhulin, 2010).

In addition to the methylation sites in the chemoreceptor signaling domain, many chemoreceptors contain an additional CheR/CheB-binding site. This consists of a C-terminal pentapeptide fused to the end of a flexible linker (Perez and Stock, 2007; Bartelli and Hazelbauer, 2011; Ortega and Krell, 2020). Pentapeptide-containing chemoreceptors have been identified in 11 different bacterial phyla (Ortega and Krell, 2020). Studies of *Escherichia coli* have shown that the affinity of CheR for the pentapeptide is 50 to 100-fold higher than the affinity for the methylation sites (Wu *et al*., 1996; Barnakov *et al*., 1999; Li and Hazelbauer, 2020). Dissociation constants of approximately 2 µM were obtained for CheR binding to the individual pentapeptide and to the pentapeptide-containing chemoreceptor (Wu *et al*., 1996), indicating that all of the determinants for high-affinity CheR binding are located in the pentapeptide. In contrast to CheR, unphosphorylated CheB bound to the pentapeptide with low affinity (Barnakov *et al*., 2002; Velando *et al*., 2020; Li *et al*., 2021). However, a stable phosphorylation mimic of *E. coli* CheB had a significantly higher affinity of about 13 µM, so that CheR and phosphorylated CheB both bind to the same high-affinity site (Li *et al*., 2021). Partial or full truncation of the C-terminal pentapeptide from chemoreceptors greatly decreases both methylation and demethylation *in vivo* and *in vitro,* preventing chemotaxis (Russo and Koshland, 1983; Yamamoto and Imae, 1993; Li *et al*., 1997; Le Moual *et al*., 1997; Okumura *et al*., 1998; Barnakov *et al*., 1999; Li and Hazelbauer, 2006; Lai *et al*., 2008; Uchida *et al*., 2022). However, many bacteria, such as *Pseudomonas putida* KT2440, lack pentapeptide containing chemoreceptors but mediate chemotactic responses that depend on the adaptation enzymes (García-Fontana *et al*., 2013; García *et al*., 2015; Martín-Mora *et al*., 2016), indicating that C-terminal pentapeptides are essential for some receptors, whereas not required for others, which corresponds to an issue that remains poorly understood.

Why are there CheR and CheB-P-binding pentapeptides at the C-terminus of chemoreceptors? It was proposed that high-affinity tethering of CheR and CheB-P to the C-terminal extension increases their local concentration, thereby enhancing their activity and leading to an optimal adaptation (Le Moual *et al*., 1997; Li and Hazelbauer, 2005; Li *et al*., 2021). In addition, we have shown previously that pentapeptides also confer specificity in their interaction with CheR and CheB in bacteria that possess multiple pathways. *Pseudomonas aeruginosa* has four chemosensory pathways, each of which contains a CheR and CheB homolog (Matilla *et al*., 2021). Of the 26 chemoreceptors, McpB (or Aer2) is the only pentapeptide-containing chemoreceptor and the sole chemoreceptor that feeds into the Che2 pathway (Ortega *et al*., 2017). Only CheR_2_ and CheB_2_, the methyltransferase and methylesterase of the Che2 pathway, but not any of the other CheR and CheB homologues bind the McpB/Aer2 pentapeptide (García-Fontana *et al*., 2014; Velando *et al*., 2020), permitting the targeting of a particular chemoreceptor with a specific CheR and CheB (García-Fontana *et al*., 2014; Velando *et al*., 2020).

All available pentapeptide sequences conserve amino acids with aromatic side chains in positions 2 and 5, whereas a significant variability is observed at the other three positions (Perez and Stock, 2007; Ortega and Krell, 2020). The inspection of the 3D structure of the CheR/pentapeptide complex of *Salmonella enterica* sv. Typhimurium shows that the aromatic side chains at positions 2 and 5 pack into two hydrophobic pockets, whereas the remaining three side chains establish hydrogen bonds and a salt bridge with CheR (Djordjevic and Stock, 1998). The conservation of amino acids at positions 2 and 5 is consistent with studies showing that mutations in these positions cause severe adaptation defects (Yamamoto and Imae, 1993; Okumura *et al*., 1998; Shiomi *et al*., 2000; Lai *et al*., 2006). Alanine-scanning mutagenesis of residues at positions 1, 3 and 4 resulted in a reduction of *in vitro* methylation and demethylation (Lai *et al*., 2006). Many bacterial strains contain a number of chemoreceptors that differ in their pentapeptide sequences (Gumerov *et al*., 2023), and the primary aim of this study consists in assessing how the naturally occurring variation of pentapeptide sequences within a bacterium impacts on CheR affinity and magnitude of chemotaxis.

We have addressed this question using *Pectobacterium atrosepticum* as a model. *P. atrosepticum* is among the top 10 plant pathogens (Mansfield *et al*., 2012) and is the causative agent of soft rot diseases in several agriculturally relevant crops (Toth *et al*., 2003). The reference strain *P. atrosepticum* SCRI1043 has 36 chemoreceptors, of which 19 possess a C-terminal pentapeptide. We have shown previously that none of these pentapeptides are recognized by unphosphorylated or the CheB-P mimic beryllium fluoride-derivatized CheB (Velando *et al*., 2020). The structure of SCRI1043 CheB can be closely superimposed onto that of *Salmonella enterica* sv. Typhimurium (Djordjevic and Stock, 1998; Velando *et al*., 2020). SCRI1043 has a single chemosensory pathway (Velando *et al*., 2020), and the chemoreceptors analyzed thus far respond to amino acids (PacB, PacC), quaternary amines (PacA), and nitrate (PacN) (Matilla *et al*., 2022b; Monteagudo-Cascales *et al*., 2023; Velando *et al*., 2023). We show here that there are significant differences in the affinity of the individual pentapeptides for CheR that are reflected in differences in the magnitude of the chemotactic responses. The differential recognition of CheR by pentapeptide may be a mechanism to bias chemotactic responses.

## Results

### SCRI1043 chemoreceptors possess 9 different pentapeptides

A sequence alignment of the C-terminal regions of all chemoreceptors of strain SCRI1043 is shown in Fig. 1A. Nineteen chemoreceptors possess a C-terminal pentapeptide that is tethered to the signaling domain via a linker sequence of 29 to 39 amino acids (Fig. 1A). In total, there were 9 different pentapeptide sequences than can be divided into three groups (Table 1). Whereas groups 1 and 2 each contained three pentapeptides that differ in a single amino acid, group 3 contained three pentapeptides with multiple differences. All peptides possess an W and an F at positions 2 and 5, respectively, but share significant variability at the remaining positions. The NWETF pentapeptide that is found in *E. coli* and *S. enterica* sv. Typhimurium receptors is present in eight receptors (Table 1, Fig. 1A). Three pentapeptides, DWTSF, NWTTF and NWEKF, were present in two different chemoreceptors, whereas the remaining pentapeptides were found in only one chemoreceptor (Table 1, Fig. 1A). SCRI1043 chemoreceptors possess different sensor domains: PAS, sCache, Cache3_Cache2, dCache, NIT, 4HB and HBM (Velando *et al*., 2020). Pentapeptides are exclusively found in chemoreceptors that contain a four-helix-bundle sensor domain, either in its monomodular (4HB) or bimodular (HBM) arrangement (Table 1). Pentapeptides NWTTF and DWTSF were found in both 4HB and HBM chemoreceptors.

**Fig. 1.**
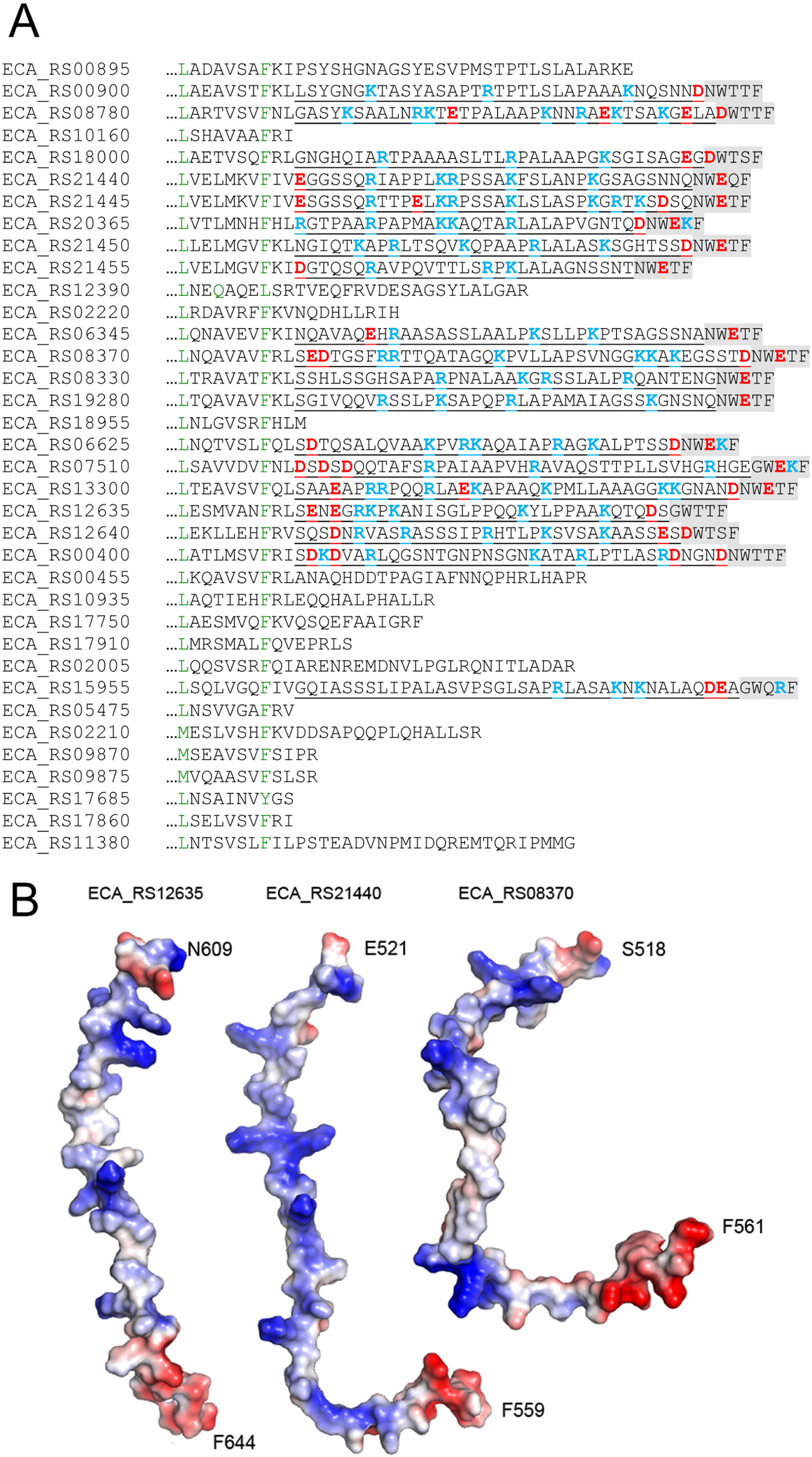
C-terminal pentapeptides at *P. atrosepticum* SCRI1043 chemoreceptors. A) The C-terminal section of the sequence alignment of SCRI1043 chemoreceptors is shown. Pentapeptides are shaded in grey. The linker sequences are underlined. Green: conserved residues in the signaling domain; red: Asp and Glu; blue: Lys and Arg. B) The electrostatic surface potential of the linker + pentapeptide sequences from three representative SCRI1043 chemoreceptors. Fragments from molecular models of the entire receptor generated by Alphafold2 (Jumper *et al*., 2021) are shown. Calculations were made using the “APBS Electrostatics” plugin of PyMOL (Schrodinger, 2010). Blue: positive charge; red: negative charge.

**Table 1.**
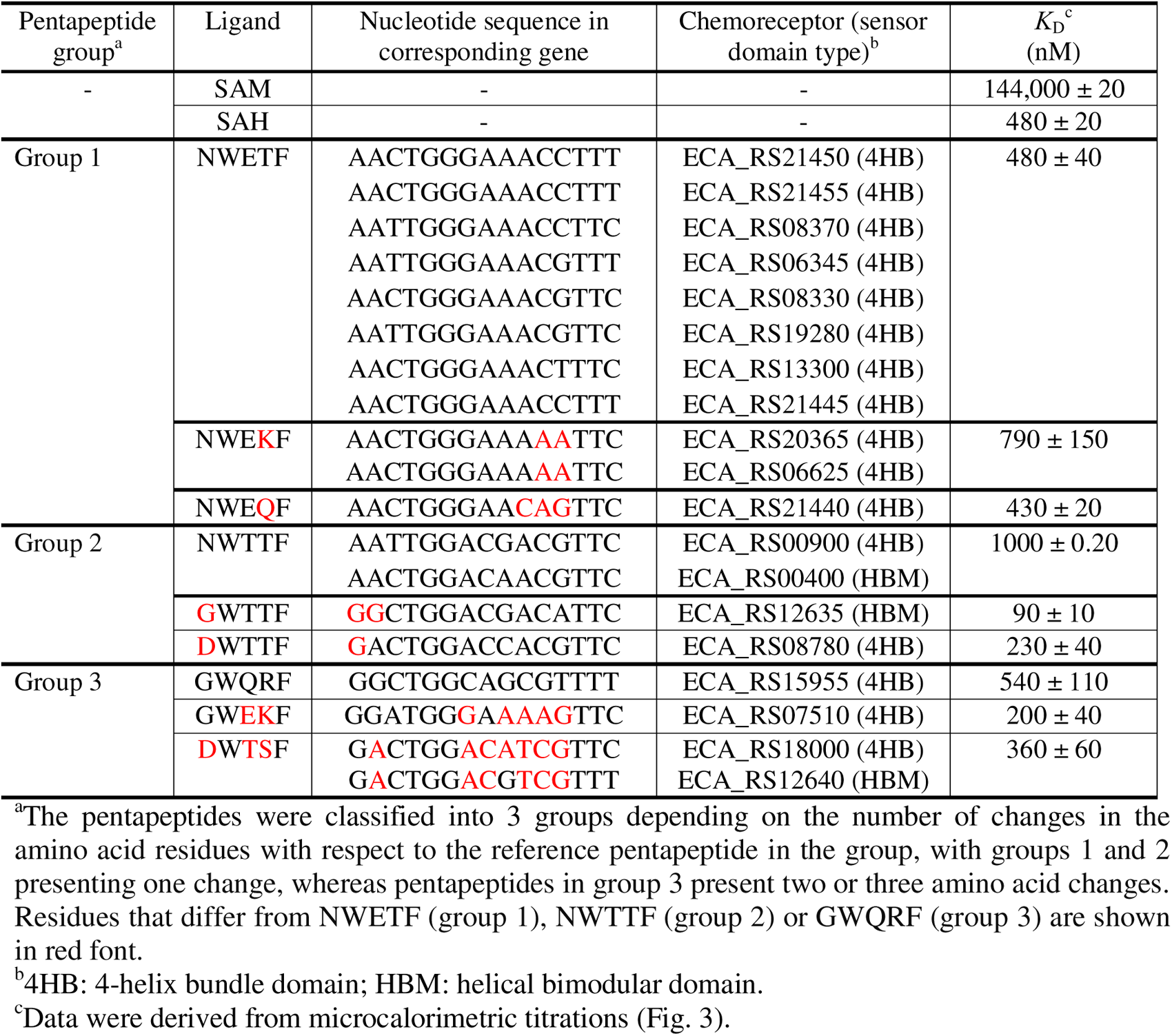
Binding parameters for the interaction of *P. atrosepticum* CheR with Sadenosylmethionine (SAM), S-adenosylhomocysteine (SAH) and pentapeptides. Amino acid changes and the corresponding alterations in the nucleotide sequence in pentapeptides from groups 1, 2 and 3 with respect to the reference pentapeptide in each group are highlighted in red color.

### Conserved charge pattern in the linker/pentapeptide sequences

So far, no conserved feature has been reported for linker sequences. They differ significantly in length, they share no apparent sequence similarities, and they are unstructured (Bartelli and Hazelbauer, 2011; Ortega and Krell, 2020). As expected, the alignment of the 19 linker sequences revealed no apparent sequence similarity (Fig. S1). However, there was a clear pattern of charge distribution (Fig. 1, Table S1). The N-terminal section of the linker frequently had a negative charge, followed by a central positively charged segment. The segment adjacent to the pentapeptide was negatively charged (Fig. 1, Table S1). The average calculated isoelectric point (pI) of the C-terminal 8 amino acids of these 19 linkers was 3.95 ± 0.41, whereas the calculated pI of the remaining linker was 10.82 ± 1.27 (Table S1). The plot of the electrostatic surface charge of Alphafold2 models of linker/pentapeptide segments from three representative chemoreceptors illustrates this pattern (Fig. 1B), which appears to be conserved. It is present in all pentapeptide-containing chemoreceptors from other chemotaxis model bacteria, including *E. coli*, *S. enterica* sv. Typhimurium, *Sinorhizobium meliloti*, *Azospirillum brasilense*, *Ralstonia solanacearum*, *Comamonas testosteroni*, and *Vibrio cholerae* (Table S2).

### SCRI1043 CheR binds to C-terminal pentapeptides with different affinities

A previous study showed that SCRI1043 CheB and a CheB-P mimic failed to bind all nine C-terminal pentapeptides present in the chemoreceptors of this strain (Velando *et al*., 2020). To investigate whether these pentapeptides bind CheR, SCRI1043 CheR was overexpressed in *E. coli* and purified from the soluble cell lysate. To verify protein integrity, microcalorimetric titrations with S-adenosylmethionine (SAM) and S-adenosylhomocysteine (SAH), the substrate and product, respectively, of the methylation reaction, were conducted (Fig. 2). Binding of SAM and SAH was characterized by *K*_D_ values of 140 ± 20 µM and 480 ± 20 nM, respectively (Table 1), revealing that the reaction product is recognized with strong preference, which agrees with data of CheR from other species (Yi and Weis, 2002; García-Fontana *et al*., 2013; García-Fontana *et al*., 2014).

**Fig. 2.**
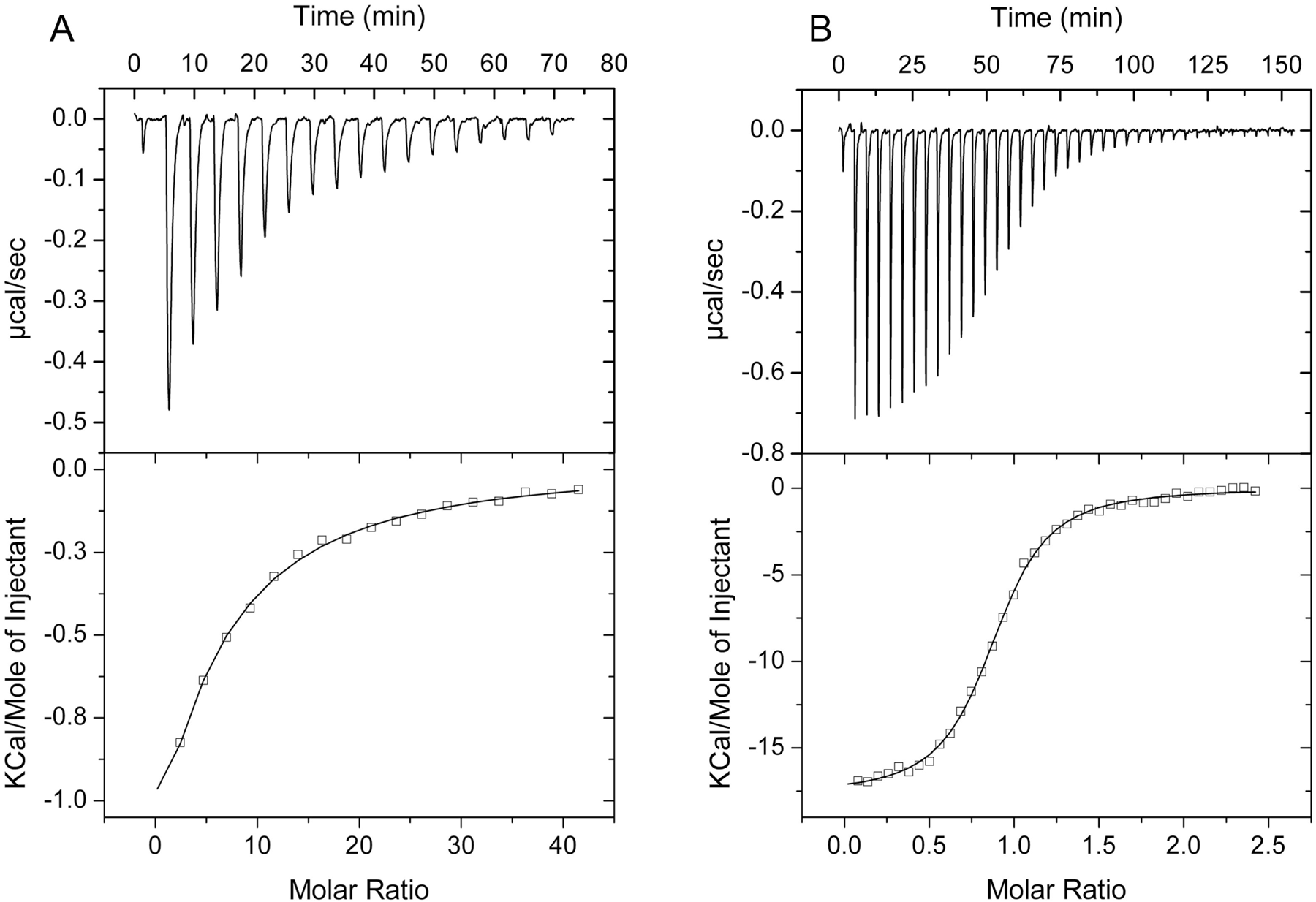
Binding of S-adenosylmethionine (SAM) and S-adenosylhomocysteine (SAH) to *P. atrosepticum* CheR. Microcalorimetric titrations of 10 µM CheR with 16 µl aliquots of 2 mM SAM (A) and of 14 µM CheR with 4.8 µl aliquots of 250 µM SAH (B). Upper panel: Raw titration data. Lower panel: Data corrected for dilution heat and concentration-normalized integrated peak areas of raw titration data. Data were fitted with the “One binding site model” of the MicroCal version of ORIGIN.

In subsequent studies, CheR was titrated with the nine different pentapeptides of SCRI1043. In all cases, peptides bound, but significant differences in affinity were observed (Fig. 3, Table 1). The titration with the NWETF pentapeptide (Fig. 3), present in *E. coli* and 8 SCRI1043 receptors, revealed a *K*_D_ value of 480 nM (Table 1), which is considerably higher affinity than *E. coli* CheR (*K*_D_=2 μM) for this peptide (Wu *et al*., 1996). NWETF derivatives containing K and Q instead of T bound with similar affinity (Table 1). Of all peptides, GWTTF bound with highest affinity (*K*_D_=90 nM).

**Fig. 3.**
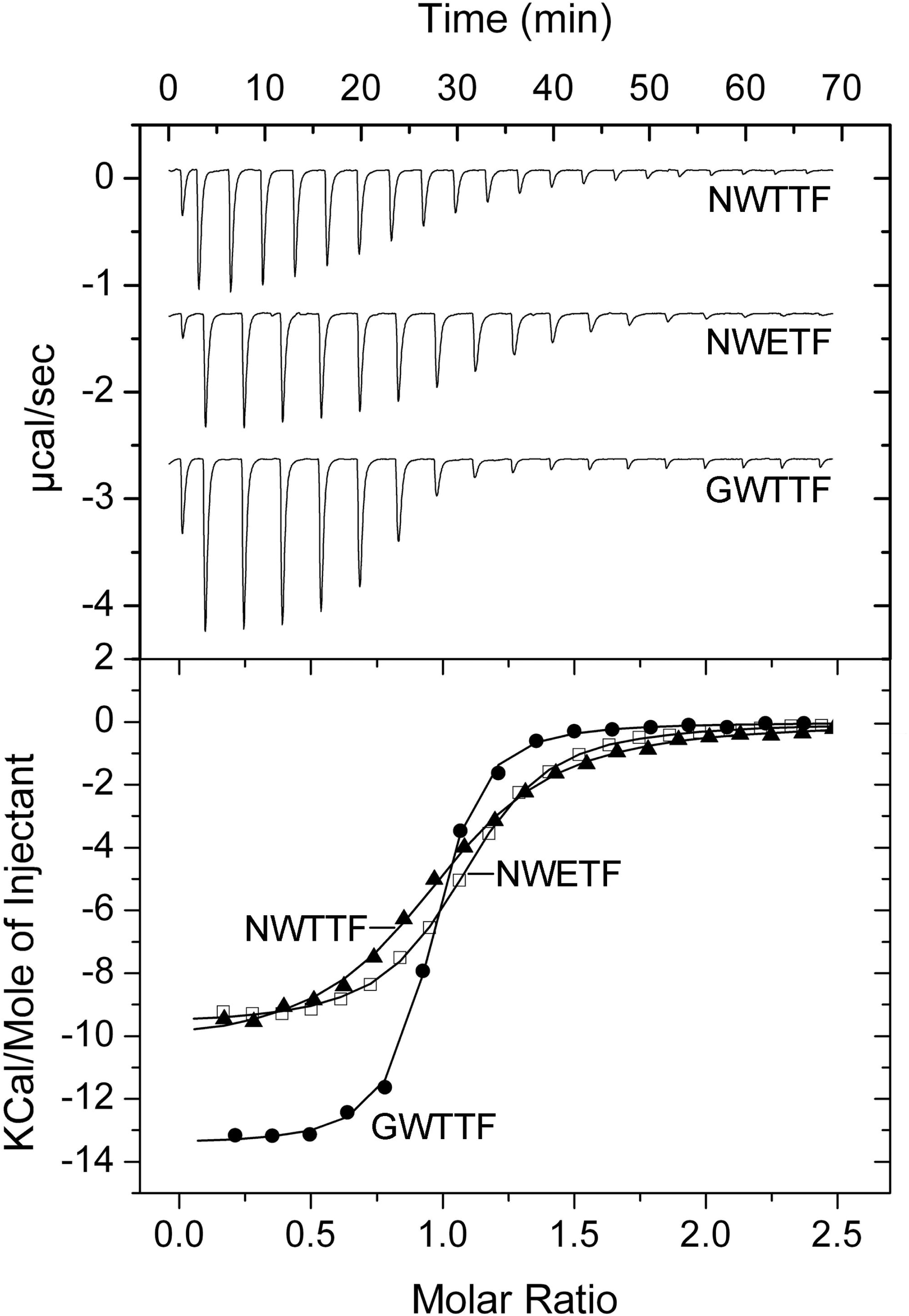
Interaction between *P. atrosepticum* pentapeptides and CheR. Upper panel: Microcalorimetric titrations of 16-20 µM CheR with 1 mM solutions of pentapeptides NWTTF, NWETF and GWTTF. Lower panel: Data corrected for dilution heat and concentration-normalized integrated peak areas of raw titration data. Data were fitted with the “One binding site model” of the MicroCal version of ORIGIN. Derived binding parameters are provided in Table 1.

Amino acid substitutions at position 1 of the pentapeptide had a pronounced effect on the binding affinity. Substitution of G by N in GWTTF resulted in the pentapeptide that bound with lowest affinity (*K*_D_=1 µM), representing a 11-fold reduction, whereas substitution with D caused a 2.6-fold decrease (Table 1). The three pentapeptides with three changes with respect to NWETF, namely GWQRF, GWEKF and DWTSF, bound with *K*_D_ values of 540, 200 and 360 nM, respectively (Table 1). Thus, variation in the residue at the less-conserved positions 1, 3 and 4 still significantly affects CheR binding affinity.

### SCRI1043 CheR is essential for chemotaxis

We have shown previously that SCRI1043 has multiple chemoreceptors for amino acids (Velando *et al*., 2023). To determine the function of CheR in SCRI1043, we constructed a *cheR* deletion mutant. Quantitative chemotaxis capillary assays showed that this mutant is unable to perform chemotaxis to casamino acids or L-aspartate. Many bacteria are strongly attracted by Krebs cycle intermediates (Matilla *et al*., 2022a). We observed a strong attraction of SCRI1043 to L-malate (Fig. 4), for which the corresponding chemoreceptor(s) remain(s) unknown. The *cheR* mutant also failed to be attracted to L-malate, indicating that CheR is essential to perform chemotaxis toward amino acids and at least one organic acid.

**Fig. 4.**
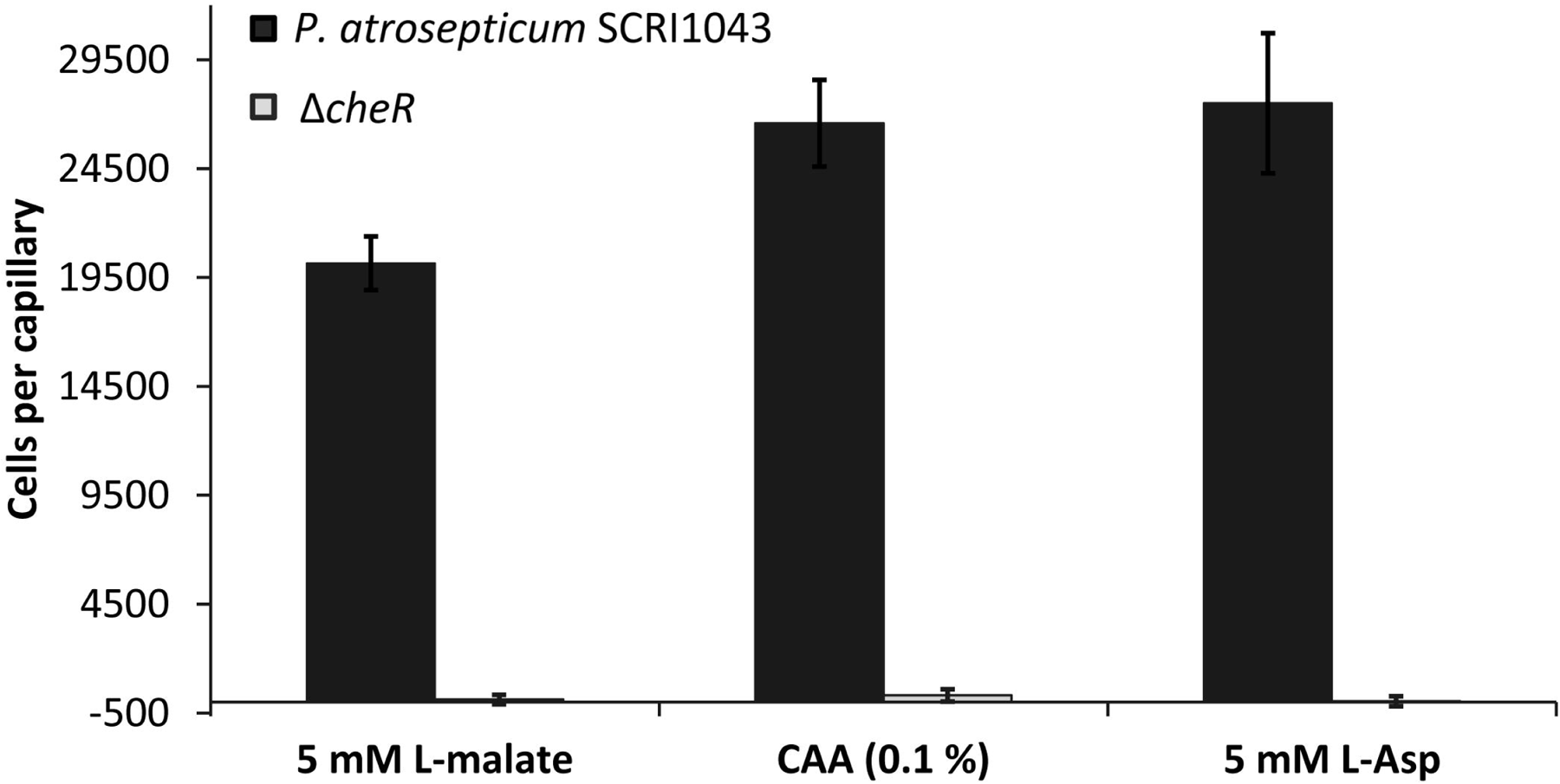
Quantitative capillary chemotaxis assays of wild-type *P. atrosepticum* SCRI1043 and a Δ*cheR* mutant towards different chemoattractants. Data have been corrected for the number of cells that swam into buffer containing capillaries (113 ± 19, 370 ± 64, and 305 ± 49 cells/capillary for L-malate, casamino acids (CAA), and L-aspartate, respectively). The means and standard deviations from three independent experiments conducted in triplicate are shown.

### Alteration of PacC mediated chemotaxis by substitution of pentapeptides

A previous study demonstrated that the strong chemotactic response of SCRI1043 to L-Asp depends entirely on the PacC chemoreceptor (Velando *et al*., 2023). As in *E. coli* Tar, PacC is encoded in the chemosensory signaling cluster and contains the NWETF pentapeptide. As shown above, CheR recognizes this pentapeptide with a *K*_D_ of 480 nM. To investigate the relevance of the CheR affinity for the pentapeptide in the chemotactic response to L-Asp, we generated two chromosomal mutations in which the region encoding the original pentapeptide was altered to encode GWTTF or NWTTF, which bind CheR with either the highest (*K*_D_=90 nM) or lowest (*K*_D_=1 µM) affinity, respectively. Control capillary chemotaxis assays showed that the responses of wt and mutant strains to casamino acids are indistinguishable (Fig. S2).

Chemotaxis experiments with the wt strain showed strong responses towards different L-Asp concentrations, confirming previously published data (Velando *et al*., 2023). Maximal responses were observed at a concentration of 1 mM. When these experiments were repeated with the mutant strain encoding PacC containing the NWTTF pentapeptide (lowest affinity), a statistically significant drop in the chemotaxis magnitude was observed at a concentrations of 1 mM L-Asp and 10 mM for the receptor containing the GWTTF pentapeptide (Fig. 5). Differences in chemotaxis mediated by the wt and mutant receptors to lower L-Asp concentrations (0.01 and 0.1 Mm) were statistically not significant (Fig. 5). These data demonstrate that the affinity of the CheR-pentapeptide interaction influences the chemotactic response. Either tighter or looser binding of CheR was associated with a decreased chemotaxis response to 1 Mm L-Asp.

**Fig. 5.**
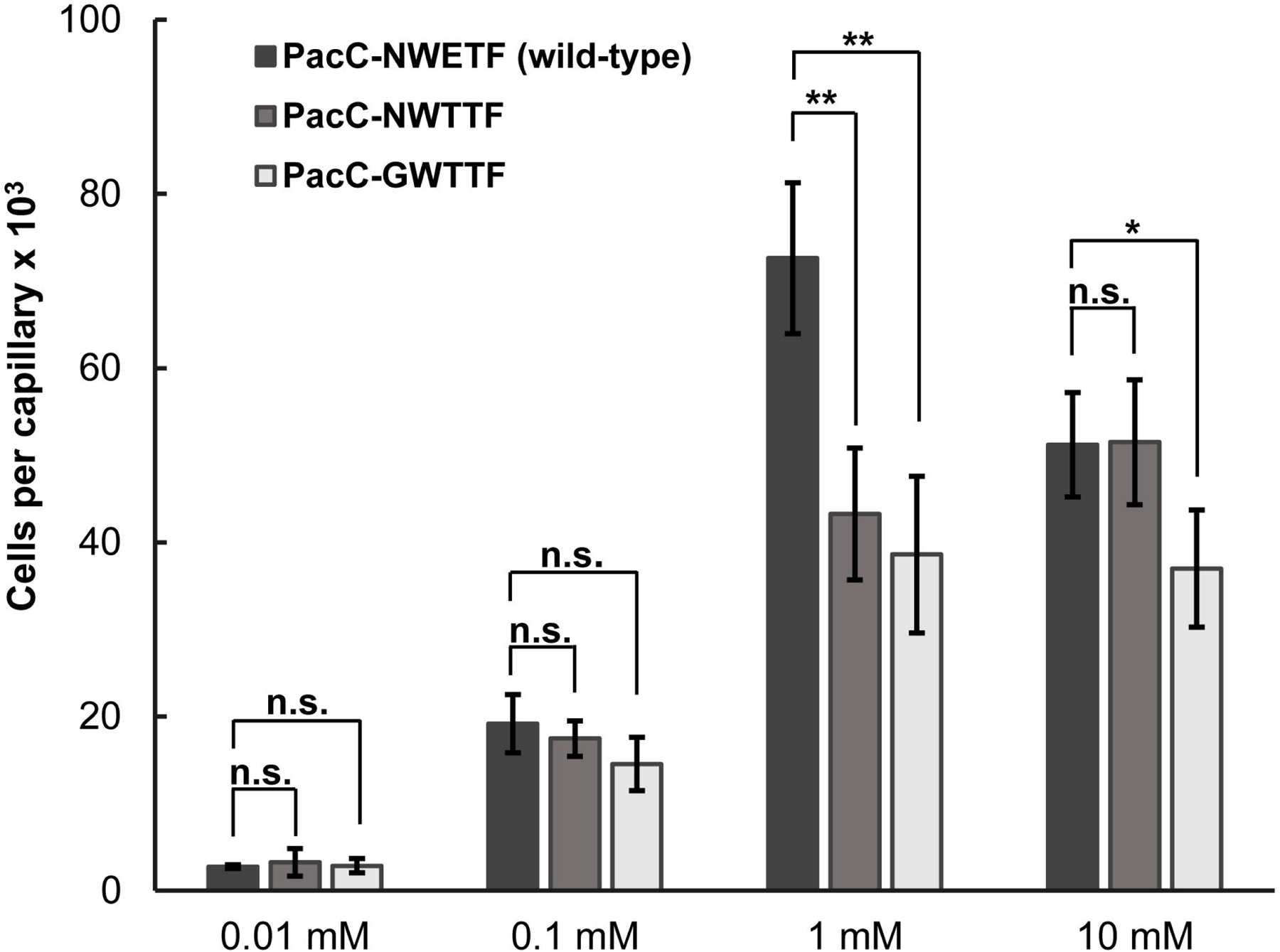
Quantitative L-Asp capillary chemotaxis assays of wild type P. atrosepticum SCRI1043 and chromosomal mutants containing variations in the PacC pentapeptide sequence. Data have been corrected for the number of bacteria that swam into buffer-containing capillaries (872 ± 248 for wt, 975 ± 517 for the PacC-NWTTF mutant, and 445 ± 109 for the PacC-GWTTF mutant). Data are the means and standard deviations from at least three biological replicates conducted in triplicate. * P<0.05 in Student’s t-test, ** P<0.01.

## Discussion

*E. coli* and *S. enterica* sv. Typhimurium are the traditional model organisms in chemotaxis research. Two of their chemoreceptors, Tar and Tsr, contain the NWETF pentapeptide and most of what we know about pentapeptide function is therefore based on the study of the NWETF pentapeptide (Wu *et al*., 1996; Djordjevic and Stock, 1998; Feng *et al*., 1999; Barnakov *et al*., 1999; Shiomi *et al*., 2000; Lai and Hazelbauer, 2005; Lai *et al*., 2006; Lai *et al*., 2008; Li and Hazelbauer, 2020). Bioinformatic studies of all available chemoreceptor sequences show that their C-terminal pentapeptides differ in sequence (Perez and Stock, 2007; Ortega and Krell, 2020), and many strains contain receptors with pentapeptides having different sequences (Gumerov *et al*., 2023). Here, we present the first study evaluating CheR recognition of the complete set of different pentapeptides within a bacterial strain.

A previous study showed that native CheB or the beryllium fluoride-modified CheB-P mimic of SCRI1043 did not interact with any of these pentapeptides (Velando *et al*., 2020), although it is unknown if an interaction with CheB-P may occur *in vivo*. In contrast, all 9 pentapeptides were recognized by SCRI1043 CheR. The affinities were between 90 nM and 1 μM, and thus higher than the affinity of *E. coli* CheR for NWETF of about 2 µM (Wu *et al*., 1996; Li and Hazelbauer, 2020), but comparable to the *K*_D_ of about 500 nM derived for the interaction of *P. aeruginosa* CheR_2_ with the GWEEF peptide of the McpB/Aer2 chemoreceptor (García-Fontana *et al*., 2014).

The importance of each of the 5 amino acids of the NWETF pentapeptide in *E. coli* has been investigated using alanine-scanning mutagenesis (Lai *et al*., 2006). Whereas mutation of W and F abolished methylation and demethylation *in vitro*, mutations of the remaining three amino acids resulted in a partial reduction of both activities. However, alanine is an amino acid that is rarely found in positions 1, 3 and 4 (Perez and Stock, 2007; Ortega and Krell, 2020). Here, we investigated naturally occurring pentapeptides and found that they differ in their affinity for CheR by up to 11-fold. These alterations are consistent with the 3D structure of the *S. enterica* CheR/pentapeptide complex showing that the N and T of the NWETF pentapeptide establish hydrogen bonds with the protein, whereas the E forms a salt bridge with a CheR Arg residue (Djordjevic and Stock, 1998). Altering these interactions by amino acid replacements might be expected to alter binding affinity.

The notion that residues at positions 1, 3, and 4 do not occur in a random fashion but are rather the result of evolution is supported by several observations. First, the sequence logo of all naturally occurring pentapeptides shows clear amino acid preference. In all three positions aspartate or glutamate residues are predominant (Ortega and Krell, 2020). Second, the single amino acid substitutions in pentapeptides of groups 1 and 2 are frequently caused by two nucleotide changes (Table 1) and are thus not single nucleotide polymorphisms.

In their natural environment, bacteria are exposed to complex mixtures of compounds, many of which are chemoeffectors. Also, in many bacteria, multiple chemoreceptors stimulate a single chemotaxis pathway (Gumerov *et al*., 2023). However, the mechanisms that permit the generation of specific chemotactic responses in the presence of complex chemoeffector mixtures is unclear.

How is the weight an individual chemoreceptor in the final response determined? One contribution is the affinity with which individual chemoeffectors are recognized by different chemoreceptors; a parameter that is known to determine the onset of a chemotactic response (Reyes-Darias *et al*., 2015; Ortega and Krell, 2020). Other factors are the level of expression and cellular abundance of chemoreceptors (Feng *et al*., 1997; Zatakia *et al*., 2018). Here, we have found that changes in the pentapeptide sequence can also alter the magnitude of the chemotactic response. Differences in affinity may result in a differential occupation of chemoreceptors with CheR. In contrast to *E. coli*, where CheB was found to bind to the pentapeptide (Li *et al*., 2021), we have failed to observe an interaction of any of the pentapeptides with *P. atrosepticum* CheB either in its unphosphorylated form or as a CheB-P mimic (Velando *et al*., 2020). However, an interaction with CheB-P may occur in the cell, and the failure to detect this interaction *in vitro* may be due to a technical issue. Considering that pentapeptides differ in their affinities for CheR, it is reasonable to suggest that such differences may also occur for binding of CheB-P. Thus, an altered interaction with CheB-P may also contribute to the dependence of PacC-mediated aspartate taxis seen with different pentapeptide sequences.

Optimal chemotaxis to L-Asp was observed when the PacC receptor contained its native NWETF pentapeptide, which has a *K*_D_ for CheR of 480 nM. When the pentapeptide was converted to NWTTF, which binds CheR with a *K*_D_ that is ∼2-fold higher (1 µM), or to GWTTF, which binds CheR with a *K*_D_ that is ∼5-fold lower (90 nM), the chemotactic responses were lower at 1 mM L-Asp. As noted above, the affinity of different pentapeptides for the other adaptation enzyme, the CheB-P demethylase, may also vary. Optimal responses were thus observed for the receptor with the naturally occurring pentapeptide, suggesting that CheR recognition at chemoreceptors has been finely tuned to achieve optimum chemotaxis.

The linker that leads to the pentapeptide does not have a conserved sequence. It is considered to be an unstructured, disordered and flexible arm that facilitates the movement of tethered adaptation enzymes within the receptor array (Bartelli and Hazelbauer, 2011). We report here that there is a conserved distribution of charge within SCRI1043 chemoreceptors (Table S1) that is also conserved in the receptors of bacteria that belong to different phylogenetic categories (Table S2). This pattern is independent of linker length: it was detected in the 29-residue linker of the *S. enterica* sv. Typhimurium Tcp chemoreceptor as well as in the 91-residue linker of *R. solanacearum* RS_RS23910 (Table S2). Our observations should prompt studies to investigate the functional relevance of this pattern. The signaling domain of chemoreceptors typically has a significant negative surface charge (Akkaladevi *et al*., 2018) (Fig. S3). The positive charge of the central linker segment may mediate a charge attraction with the signaling domain, whereas the negative charge of the C-terminal portion of the linker that connects to the pentapeptide may cause a repulsion away from the signaling domain. The notion that this conserved charge pattern is of functional relevance is supported by a study of *E. coli* Tar (Lai *et al*., 2008). The authors generated alanine-substitution mutants of the lysine and three arginine residues present in the Tar linker (Table S2). Whereas receptors with individual substitutions supported chemotaxis equivalent to that supported by wild-type Tar, combining all four changes decreased chemotaxis by about 50%. Importantly, the singly substituted receptors showed a significant increase in CheA activity that was increased when multiple substitutions were combined. The authors concluded that the Ala substitutions either stabilize the ON state or destabilize the OFF state. Whereas the initial 30 amino acids of this linker have a calculated pI of 10.74 (Table S2), replacing all four positively charged amino acids with Ala lowered the pI to 3.80. It was suggested that the positively charged linker either interacts with the negatively charged polar head groups of phospholipids or with charged residues in the signaling domain of the receptor (Lai *et al*., 2008). Combined with the observation that this charge pattern is conserved, this result supports the idea that this organization of the linker is of fundamental functional relevance.

Collectively, we show here that naturally occurring variations in the pentapeptide sequence cause important alterations in the binding affinities of CheR. Maximal responses were obtained for a chemoreceptor containing its naturally occurring pentapeptide, suggesting that the pentapeptide sequence has been fine-tuned to guarantee maximal responses. A conserved charge pattern was observed in linker sequences that is likely to contribute to the spatial organization of the linker within the chemoreceptor array. Our study forms the basis for investigations on how pentapeptide sequence variation affects the magnitude of chemotaxis in other bacteria and to determine the functional relevance of the observed charge pattern of the linker.

### Experimental procedures

*Bacterial strains and growth conditions*: Bacterial strains used in this study are listed in Table S3. SCRI1043 and its derivative strains were routinely grown at 30 °C in Luria Broth (5 g/l yeast extract, 10 g/l bacto tryptone, 5 g/l NaCl) or minimal medium (0.41 mM MgSO_4_, 7.56 mM (NH_4_)_2_SO_4_, 40 mM K_2_HPO_4_, 15 mM KH_2_PO_4_) supplemented with 0.2% (w/v) glucose as carbon source. *E. coli* strains were grown at 37 °C in LB. *Escherichia coli* DH5α was used as a host for gene cloning. Media for propagation of *E. coli* β2163 were supplemented with 300 mM 2,6-diaminopimelic acid. When appropriate, antibiotics were used at the following final concentrations (in μg ml^−1^): kanamycin, 50; streptomycin, 50; ampicillin, 100. Sucrose was added to a final concentration of 10% (w/v) when required to select derivatives that had undergone a second cross-over event during marker exchange mutagenesis.

*In vitro nucleic acid techniques:* Plasmid DNA was isolated using the NZY-Tech miniprep kit. For DNA digestion, the manufacturer’s instructions were followed (New England Biolabs and Roche). Separated DNA fragments were recovered from agarose using the Qiagen gel extraction kit. Ligation reactions were performed as described in (Sambrook *et al*., 1989). PCR reactions were purified using the Qiagen PCR Clean-up kit. PCR fragments were verified by DNA sequencing that was carried out at the Institute of Parasitology and Biomedicine Lopez-Neyra (CSIC; Granada, Spain). Phusion^®^ high fidelity DNA polymerase (Thermo Fisher Scientific) was used in the amplification of PCR fragments for cloning.

*CheR overexpression and purification*: The DNA sequence encoding CheR of SCRI1043 (ECA_RS08375) was amplified by PCR using the oligonucleotides indicated in Table S4 and genomic DNA as template. The PCR product was digested with NheI and SalI and cloned into pET28b(+) linearized with the same enzymes. The resulting plasmid pET28b-CheR was verified by DNA sequencing and transformed into *E. coli* BL21-AI^TM^. The strain was grown under continuous stirring (200 rpm) at 30 °C in 2-liter Erlenmeyer flasks containing 500 ml of LB medium supplemented with 50 μg/ml kanamycin. At an OD_660_ of 0.6, protein expression was induced by the addition of 0.1 mM isopropyl β-D-1-thiogalactopyranoside (IPTG) and 0.2 % (w/v) L-arabinose. Growth was continued at 16 °C overnight prior to cell harvest by centrifugation at 10 000 x *g* for 30 min. The cell pellet was resuspended in buffer A (40 mM KH_2_PO_4_*/*K_2_HPO_4_, 10 mM imidazole, 10% (v/v) glycerol, 1 mM β-mercaptoethanol, pH 7.5) and cells were broken by French press treatment at 62.5 lb/in^2^. After centrifugation at 20 000 x *g* for 30 min, the supernatant was loaded onto a 5-ml HisTrap HP columns (Amersham Biosciences) equilibrated with buffer A and eluted with an imidazole gradient of 40–500 mM in the same buffer. Protein-containing fractions were pooled, and dialyzed overnight against 40 mM KH_2_PO_4_*/*K_2_HPO_4_, 10% (v/v) glycerol, 1 mM β-mercaptoethanol, pH 7.0. All manipulations were carried out at 4 °C. Experiments were conducted with the hexa-histidine tagged protein.

*Isothermal titration calorimetry (ITC):* All experiments were conducted on a VP-microcalorimeter (MicroCal, Amherst, MA) at 25 °C. CheR was dialyzed overnight against 40 mM KH_2_PO_4_*/*K_2_HPO_4_, 10% (v/v) glycerol, 1 mM β-mercaptoethanol, pH 7.0, adjusted to a concentration of 10–20 μM, placed into the sample cell and titrated with 3.2–4.8 μl aliquots of 0.1–1 mM peptide solutions (purchased from GenScript, Piscataway, NJ, USA), 2 mM SAM or 250 µM SAH. All solutions were prepared in dialysis buffer immediately before use. The mean enthalpies measured from the injection of the peptide into the buffer were subtracted from raw titration data prior to data analysis with the MicroCal version of ORIGIN. Data were fitted with the ‘One binding site model’ of ORIGIN.

*Construction of a mutant deficient in* cheR. A chromosomal *cheR* mutant of SCRI1043 was constructed by homologous recombination using a derivative plasmid of the suicide vector pKNG101. The up- and downstream flanking regions of *cheR* were amplified by PCR using the oligonucleotides listed in Table S4, which were subsequently digested with EcoRI/BamHI and BamHI/PstI, respectively, and ligated in a three-way ligation into the EcoRI/PstI sites of pUC18Not to generate pUC18Not_ΔcheR. Subsequently, a 0.95-kb BamHI fragment containing the Km3 cassette of p34S-Km3 was inserted into the BamHI site of pUC18Not_ΔcheR, giving rise to pUC18Not_ΔcheR_km3. Lastly, a ∼2.5-kb NotI fragment of pUC18Not_ΔcheR_km3 was cloned at the same site into pKNG101, resulting in pKNG101_ΔcheR. This plasmid was transferred to SCRI1043 by biparental conjugation using *E. coli* β2163. Then, cells were spread onto LB plates containing kanamycin at 50 µg/ml. Selected merodiploid colonies were spread onto LB plates containing 10 % (w/v) sucrose to select derivatives that had undergone a second cross-over event during marker exchange mutagenesis. All plasmids and the final mutant were confirmed by PCR and sequencing.

*Construction of SCRI1043 mutant strains* pacC*-NWTTF and* pacC*-GWTTF*. Chromosomal mutants of SCRI1043 encoding PacC variants with altered pentapeptides were constructed by homologous recombination using derivative plasmids of the suicide vector pKNG101. Briefly, an overlapping PCR mutagenesis approach was employed to construct *pacC* gene variants in which the region encoding the original NWETF pentapeptide sequence was altered to encode GWTTF or NWTTF, using oligonucleotides listed in Table S4. The corresponding ∼1-kb PCR products were then digested with EcoRI/PstI and cloned into the same sites of pUC18Not to generate pUC18Not-pacC-NWTTF and pUC18Not-pacC-GWTTF. Subsequently, ∼1.1-kb NotI fragments of these plasmids were cloned at the same site in pKNG101 to generate pKNG-pacC-NWTTF and pKNG-pacC-GWTTF, which were transferred to *P. atrosepticum* SCRI1043 by biparental conjugation using *E. coli* β2163. After overnight incubation at 30 °C, cells were spread onto LB plates containing streptomycin at 50 µg/ml. Selected merodiploid colonies were spread onto LB plates containing 10 % (w/v) sucrose to select derivatives that had undergone a second cross-over event during marker exchange mutagenesis. All plasmids and final mutants were confirmed by PCR and sequencing.

*Quantitative capillarity chemotaxis assays*. Overnight bacterial cultures of SCRI1043 were grown at 30 °C in minimal medium. At an OD_660_ of 0.3-0.4, the cultures were washed twice by centrifugation (1,667[×[*g* for 5 min at room temperature) and subsequent resuspension in chemotaxis buffer (50 mM K_2_HPO_4_/KH_2_PO_4_, 20 μM EDTA and 0.05% (v/v) glycerol, pH 7.0). Cells were diluted to an OD_660_ of 0.1 in the same buffer. Subsequently, 230 μl aliquots of the resulting bacterial suspension were place into 96-well plates. One-microliter capillary tubes (P1424, Microcaps; Drummond Scientific) were heat-sealed at one end and filled with either the chemotaxis buffer (negative control) or chemotaxis buffer containing the chemoeffectors to test. The capillaries were immersed into the bacterial suspensions at its open end. After 30 min at room temperature, the capillaries were removed from the bacterial suspensions, rinsed with sterile water and the content expelled into tubes containing 1 ml of minimal medium salts. Serial dilutions were plated onto minimal medium supplemented with 15 mM glucose as carbon source. The number of colony forming units was determined after 36 h incubation at 30 °C. In all cases, data were corrected with the number of cells that swam into buffer containing capillaries. Data are the means and standard deviations from at least three biological replicates conducted in triplicate.

## Acknowledgements

We are indebted to Dr. Michael Manson for his constructive scientific criticism and editing the English of this manuscript. We would also like to thank Dr. Gerald Hazelbauer for his advice. This work was supported by the Spanish Ministry for Science and Innovation/*Agencia Estatal de Investigación* 10.13039/501100011033 through grants PID2020-112612GB-I00 to TK and PID2019-103972GA-I00 to MAM, CSIC grant 2023AEP002 to MAM, and the Junta de Andalucía (grant P18-FR-1621 to TK).

## Author contributions

FV: investigation, formal analysis, visualization, writing – review & editing; EMC: investigation, formal analysis, visualization, writing – review & editing; MAV: Conceptualization, funding acquisition, project administration, supervision, writing – review & editing; TK: Conceptualization, funding acquisition, project administration, supervision, writing – original draft preparation, writing – review & editing

## Abbreviated Summary

The nine different pentapeptides present in *P. atrosepticum* SCRI1043 differ in their affinity for the CheR methyltransferase up to 11-fold. Replacement of the naturally occurring NWETF pentapeptide at the PacC chemoreceptor with pentapeptides the bind with lower and higher affinity to CheR reduces the magnitude of chemotaxis to 1 mM L-Asp. The linker sequences showed a charge pattern that was conserved in all pentapeptide containing chemoreceptors of other chemotaxis model strains.

## Abbreviations

4HB: four-helix bundle domain
HBM: helical bimodular domain
ITC: isothermal titration calorimetry
LBD: ligand binding domain
SCRI1043: *Pectobacterium atrosepticum* SCRI1043
SAM: S-adenosylmethionine
SAH: S-adenosylhomocysteine.

