## Supplementary Figures and Tables for "Differential CheR affinity for chemoreceptor C-terminal pentapeptides biases chemotactic responses"

to

by

Félix Velando, Elizabet Monteagudo-Cascales, Miguel A. Matilla and Tino Krell

```
ECA_RS00900      LSYGNGKTA-SYASAPTRTPTL-S-LAP--AAAKNQSNND--
ECA_RS08780      ---GASYKS-AALNRKTETPAL-A-APKNNRAEKTSAKGELA
ECA_RS20365      -GTPAARPA-P-MAKKAQTAR----LALAPVG----NTQD--
ECA_RS13300      AAEAPRRPQ-QRLAEKAPAAQK-P-MLLAAAGGKKGNAND--
ECA_RS21440      ---GGSSQ---RIAPPLKRPSS-AKFSLANPKGSAGSNNQ--
ECA_RS21445      --ESGSSQ---RTTPELKRPSS-AKLSLASPKGRTKSDSQ--
ECA_RS21450      ---GIQTKA-PRLTSQVKQPAA-PRLALASKSGHTSSD----
ECA_RS21455      ---GTQSQ---RAVPQVTTLSR-PKLALAGNSSNT-------
ECA_RS18000      ---GNGHQI-ARTPAAAASLTLRPALAAPGKSGISAGEG---
ECA_RS19280      ---GIVQQVRSSLPKSAPQPRLAPAMAIAGSS--KGNSNQ--
ECA_RS15955      --QIASSSLIPALASVPSGLSA-PRLASAKNKNALAQDEA--
ECA_RS08370      EDTGSFRR--TTQATAGQKPVLLAPSVNGGKKAKEGSSTD--
ECA_RS06345      --QAVAQEH--RAASASSLAAL-PKSLLPKPTS-AGSSNA--
ECA_RS08330      SHLSSGHSA-PARPNALAAKGR-SSLALPRQAN---TENG--
ECA_RS12640      ---SQSDN---RVASRASSSI--PRHTLPKSVSAKAASSES-
ECA_RS06625      ----DTQS---ALQVAAKPVRK-AQAIAPRAGKALPTSSD--
ECA_RS07510      -SDSDQQTAFSRPAIAAPVHRAVAQSTTPLLS-VHGRHGE--
ECA_RS12635      --ENEGRK--PKANISGLPPQ--QKYLPPAAK---QTQDS--
ECA_RS00400      -DKDVARLQ--GSNTGNPNSGNKATARLPTLAS-RDNGND--
```

**Fig. S1) Alignment of the linker sequences of the 19 chemoreceptors from *Pectobacterium atrosepticum* SCRI1043 containing C-terminal pentapeptides**. The alignment was done using the CLUSTALW algorithm of the npsa suite (Combet *et al.*, 2000) using the GONNET weight matrix, a gap opening penalty of 10 and a gap extension penalty of 0.2.

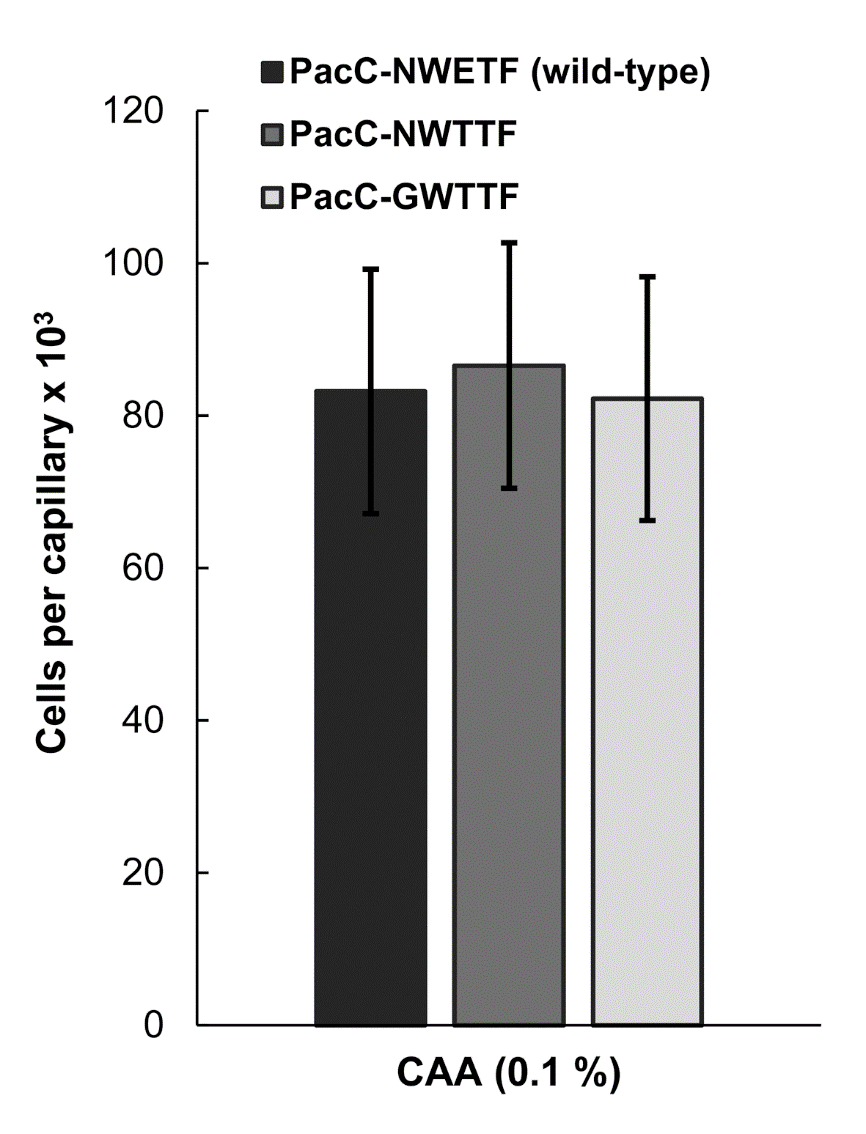

**Fig. S2) Quantitative capillary chemotaxis assays of *P. atrosepticum* SCRI1043 and mutants PacC-NWTTF and PacC-GWTTF to 0.1 % (w/v) casamino acids (CAA).** Data have been corrected with the number of bacteria that swam into buffer containing capillaries (908 ± 270 for wild-type strain, 975 ± 198 for PacC-NWTTF and 467 ± 302 for PacC-GWTTF). Data are the means and standard deviations of at least three biological conducted in triplicate.

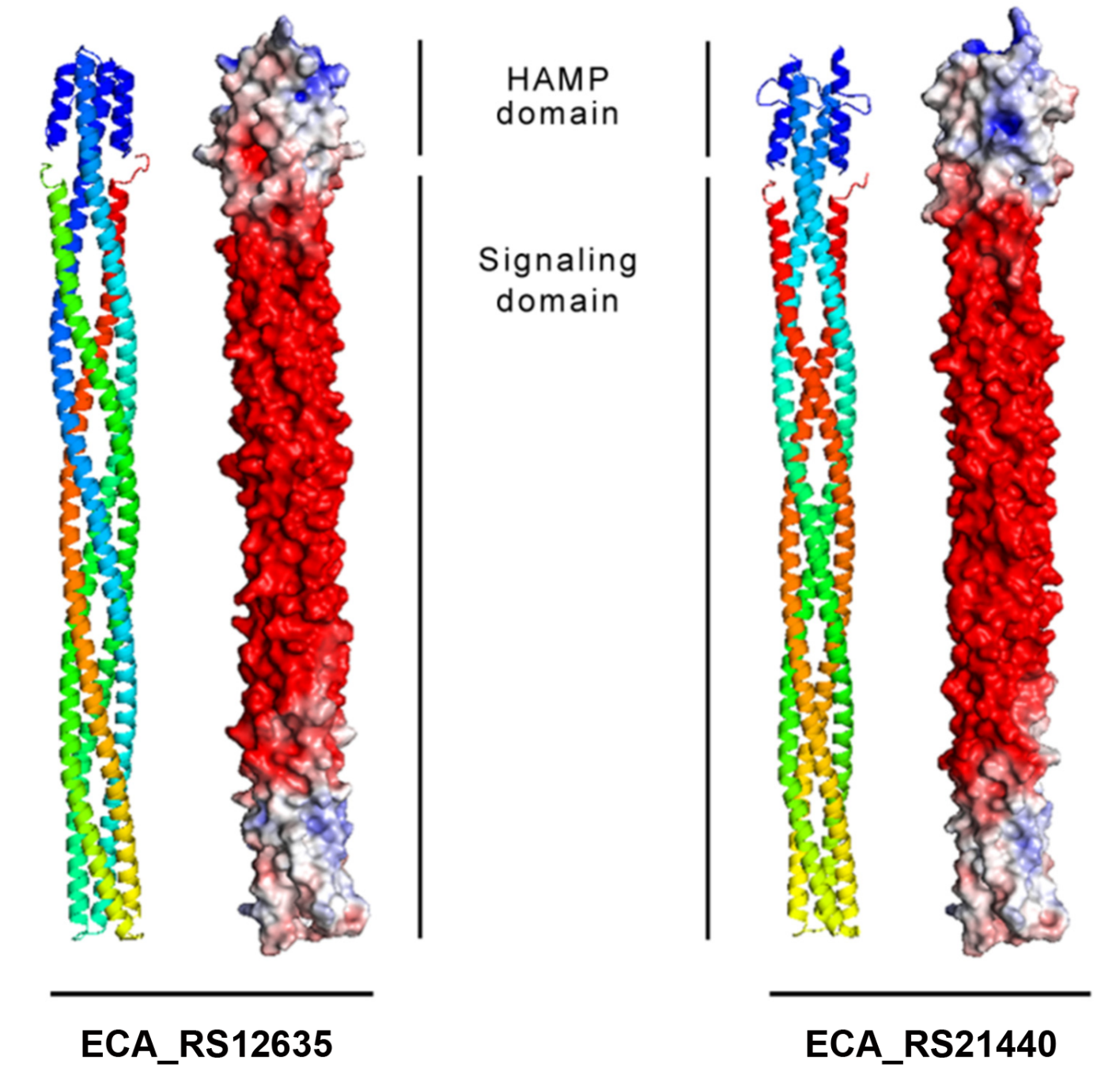

**Fig. S3) Charge distribution at the cytosolic fragments of *P. atrosepticum* chemoreceptors.** Homology models of the cytosolic fragments of the ECA_RS12635 and ECA_RS21440 receptors were produced using SwissModel (Waterhouse *et al.*, 2018). Ribbon diagrams are shown on the left and surface charges as calculated by PyMol (The PyMOL Molecular Graphics System, Version 1.3, Schrödinger, LLC., n.d.) are shown on the right. Red: negative charge, blue: positive charge. Two representative chemoreceptors are shown.

**Table S1) The sequence-derived isoelectric points (pI) of different sections of the linker/pentapeptide sequences of the 19 pentapeptide containing chemoreceptors of *P. atrosepticum*.**

| **Receptor** | **Linker + pentapeptide sequence^a^** | **Theoretical pI^b^** | |
| --- | --- | --- | --- |
|  |  | **C-terminal 8 amino acids** | **Remaining linker** |
| ECA_RS00900 | LSYGNGKTASYASAPTRTPTLSLAPAAAKNQSNNDNWTTF | 3.80 | 10.00 |
| ECA_RS08780 | GASYKSAALNRKTETPALAAPKNNRAEKTSAKGELADWTTF | 3.67 | 10.29 |
| ECA_RS18000 | GNGHQIARTPAAAASLTLRPALAAPGKSGISAGEGDWTSF | 3.67 | 12.01 |
| ECA_RS21440 | EGGSSQRIAPPLKRPSSAKFSLANPKGSAGSNNQNWEQF | 4.00 | 11.10 |
| ECA_RS21445 | ESGSSQRTTPELKRPSSAKLSLASPKGRTKSDSQNWETF | 3.67 | 11.07 |
| ECA_RS20365 | RGTPAARPAPMAKKAQTARLALAPVGNTQDNWEKF | 4.37 | 12.31 |
| ECA_RS21450 | NGIQTKAPRLTSQVKQPAAPRLALASKSGHTSSDNWETF | 3.67 | 12.02 |
| ECA_RS21455 | DGTQSQRAVPQVTTLSRPKLALAGNSSNTNWETF | 4.00 | 10.84 |
| ECA_RS06345 | NQAVAQEHRAASASSLAALPKSLLPKPTSAGSSNANWETF | 4.00 | 9.99 |
| ECA_RS08370 | SEDTGSFRRTTQATAGQKPVLLAPSVNGGKKAKEGSSTDNWETF | 3.67 | 9.99 |
| ECA_RS08330 | SSHLSSGHSAPARPNALAAKGRSSLALPRQANTENGNWETF | 4.00 | 12.30 |
| ECA_RS19280 | SGIVQQVRSSLPKSAPQPRLAPAMAIAGSSKGNSNQNWETF | 4.00 | 12.02 |
| ECA_RS06625 | SDTQSALQVAAKPVRKAQAIAPRAGKALPTSSDNWEKF | 3.67 | 11.10 |
| ECA_RS07510 | DSDSDQQTAFSRPAIAAPVHRAVAQSTTPLLSVHGRHGEGWEKF | 5.40 | 6.92 |
| ECA_RS13300 | SAAEAPRRPQQRLAEKAPAAQKPMLLAAAGGKKGNANDNWETF | 3.67 | 11.07 |
| ECA_RS12635 | SENEGRKPKANISGLPPQQKYLPPAAKQTQDSGWTTF | 4.00 | 9.82 |
| ECA_RS12640 | SQSDNRVASRASSSIPRHTLPKSVSAKAASSESDWTSF | 3.67 | 11.72 |
| ECA_RS00400 | SDKDVARLQGSNTGNPNSGNKATARLPTLASRDNGNDNWTTF | 3.80 | 9.98 |
| ECA_RS15955 | GQIASSSLIPALASVPSGLSAPRLASAKNKNALAQDEAGWQRF | 4.37 | 11.17 |
| **Average** | | **3.95 ± 0.41** | **10.82 ± 1.27** |

^a^Positively and negatively charged amino acids are highlighted in blue and red, respectively. The sequences corresponding to the pentapeptides are boxed.

^b^Values were determined using the ProtParam algorithm (<https://web.expasy.org/cgi-bin/protparam/protparam>).

**Table S2) The sequence-derived isoelectric points (pI) of different sections of the linker/pentapeptide sequences of chemoreceptors from a selection of model strains used for chemotaxis research.**

| **Strain/**  **Phylogenetic class** | **Receptor** | **Linker + pentapeptide sequence^a^** | **Theoretical pI^b^** | |
| --- | --- | --- | --- | --- |
|  |  |  | **C-terminal 8 amino acids** | **Remaining linker** |
| *Sinorhizobium meliloti* RU11/001 (α-proteobacteria) | McpT | VEREAVRTVVPAKDASRPVASPARRMMGTVARAFGNGSAAVARDDWEEF | 3.92 | 11.71 |
|  | McpX | IAPQAAQAQASAEMLRGTAERMRAAAPAENRPAQAPRSAAYSNSTQRVLAKTSGANALAQDNWEEF | 3.57 | 11.34 |
|  | McpY | GRDASPASDNRMEAPHSPTRLHATAKTLRSGTRSNLALAPAADDWENF | 3.49 | 11.42 |
|  | McpW | HGQARETAIRTAAIQPPGRAGSATASAPARAATATAAAPVSLRKPAAAQSSTRPAPSPAKALMGKLAGAFGNRPGSTPSVTASGENWEEF | 3.67 | 12.40 |
| Azospirillum brasilense Sp7 (α-proteobacteria) | AMK58_RS04195 | FFKLDAAAFMAAPAGHQGGGAPALRRSIAPTKPAPAKPAARKAAPPARPAAGKAASATLQRASADEDDWKEF | 3.77 | 12.19 |
|  | AMK58_RS20975 | KLDPRAMPVAVPVAVATTAAAPRHVAPVVRPTRAAAKPSARAVSGAALRRNVSEDDDWKEF | 3.77 | 12.40 |
| *Ralstonia solanacearum* GMI1000 (β-proteobacteria) | RS_RS05755 | TMSAVRSVVAKAEPTIAPPVPKAAEAGPATPVPVAAPVAAPAQAPRREPVKPSPEAARPKLQPRKAPVLAKRAARPDTAPATVPAASPKPLHQPLVAAGDDADWETF | 3.37 | 11.36 |
|  | RS_RS05760 | LDETDGVVAAAASKPVRTAPAAAVKPAAKPPRPVVRKRGPSPKAVAPATVPVVAPAAAPVVAPVKAQPLAAVAGSDGDWETF | 3.49 | 11.10 |
|  | RS_RS19595 | QGGGEVRAAAPAVRAVAAAKPAAAATVARPAVRKPAATVRARAPRKPAAEAPVAKAASSPAPAPAEPAGGKLALSAAVDQDDWTQF | 3.42 | 11.89 |
|  | RS_RS23910 | TRADAPAQTVARPARPAAAPVKAPARATALARRPAPKAAVARPPATAGEATPSATASSNEQQRTRQAAAKAVVPAPKKQAPKLVAAGGDESDWETF | 3.43 | 12.11 |
| *Comamonas thiooxydans* (formely *Comamonas testosteron*i) CNB-2 (β-proteobacteria) | CTCNB1_RS04260 | KLGRNAEAFTTVSAVLPKPLATIKALNPAAQPAVVAPRPVASRSKEQSLASTAEDDSWEQF | 3.43 | 11.07 |
|  | CTCNB1_RS08490 | FQLGNQAQSFAAAALKNRVEPVAKAGMSAGPRAAALAGQGPGKAVPAPVKTAQPALSNDQDWEQF | 3.49 | 11.17 |
|  | CTCNB1_RS08495 | LPASALYDQAPKAAARAAAPAVVAAAAPARQAPVRRVAARPAPMPAAPRSLMGGAVAGESATARNVQKSSLDEDDWETF | 3.33 | 11.88 |
|  | CTCNB1_RS10330 | TLSEQIQGPDSGMPTTSASVTKGTLAPNLSKPLQASLASIKSTVERRPASIEGTQRSTGKTDVDWETF | 3.49 | 9.99 |
|  | CTCNB1_RS15035 | LSEHEQHASAAMQAASVNPTKTATVAKVSESPPKPLMATKSPVLRRPVAIAGTQQATNKNGDDWETF | 3.49 | 10.17 |
|  | CTCNB1_RS15335 | LGAQSGLQATPALAPAALKSAPARAAIKAAAPLKAPSAALKRPAMSVGSKPPVLSPAKSAAAPQPQKAATADDGEWESF | 3.43 | 12.05 |
| *Escherichia coli* K12  (γ-proteobacteria) | Tar | FRLAASPLTNKPQTPSRPASEQPPAQPRLRIAEQDPNWETF | 3.67 | 11.54 |
|  | Tsr | FRIQQQQRETSAVVKTVTPAAPRKMAVADSEENWETF | 3.67 | 10.90 |
| *Salmonella enterica* sv. Typhimurium LT2 (γ- proteobacteria) | Tar | FRLASRPLAVNKPEMRLSVNAQSGNTPQSLAARDDANWETF | 3.49 | 12.00 |
|  | Tsr | IHQQQQRAREVAAVKTPAAVSSPKAAVADGSDNWETF | 3.67 | 9.99 |
|  | McpB | FRLSASEPQQRVTAKAAPGVQRMASAPAQSTDEWVSF | 3.67 | 11.71 |
|  | Tcp | FRIQKQPRREASPTTLSKGLTPQPAAEQANWESF | 3.80 | 11.72 |
| *V. cholerae* O1 biovar El Tor N16961  (γ-proteobacteria) | VC0098 | DQDTSAPSLLKAVNKRPQSAPVTRHPASHIAKTPAKITAKASSRAQPVMQVAHDEEWESF | 4.00 | 11.75 |
| **Average** | | | **3.59 ± 0.18** | **11.48 ± 0.63** |

^a^Positively and negatively charged amino acids are highlighted in blue and red, respectively. The sequences corresponding to the pentapeptides are boxed.

^b^Values were determined using the ProtParam algorithm (<https://web.expasy.org/cgi-bin/protparam/protparam>).

**Table S3) Strains and plasmids used in this study.**

| **Strains and plasmids** | **Genotype or relevant characteristics^a^** | **Reference** |
| --- | --- | --- |
| **Strains** | | |
| *Escherichia coli* BL21-AI | F- *ompT hsdS*_B_ (r_B_^-^m_B_^-^) *gal dcm araB*::*T7RNAP-tetA* | Invitrogen |
| *E. coli* DH5α | F^–^ *endA1* *glnV44* *thi-1*  *recA1*  *relA1*  *gyrA96 deoR* *nupG* *purB20* φ80d*lacZ*ΔM15 Δ(*lacZYA-argF*)U169, hsdR17(*r_K_*^–^*m_K_*^+^), λ^–^ | (Woodcock *et al.*, 1989) |
| *E. coli* β2163 | F- RP4-2-Tc::Mu Δ*dapA*::(*erm-pir*); Km^R^, Em^R^ | (Demarre *et al.*, 2005) |
| *Pectobacterium atrosepticum* SCRI1043 | Wild type strain | (Bell *et al.*, 2004) |
| *P. atrosepticum* SCRI1043 Δ*cheR* | SCRI1043 Δ*cheR*::km. Constructed by marker exchange; Km^R^ | This study |
| *P. atrosepticum* SCRI1043 *pacC*-NWTTF | SCRI1043 *pacC* mutant in the chromosome in which the region encoding the original pentapeptide sequence was altered to encode NWTTF | This study |
| *P. atrosepticum* SCRI1043 *pacC*-GWTTF | SCRI1043 *pacC* mutant in the chromosome in which the region encoding the original pentapeptide sequence was altered to encode GWTTF | This study |
| **Plasmids** | | |
| pET28b(+) | Km^R^; Protein expression plasmid | Novagen |
| pET28b-CheR | Km^R^; pET28b(+) derivative containing *P. atrosepticum* *cheR* (*ECA_RS08375*) | This study |
| pUC18Not | Ap^R^; identical to pUC18 but with two NotI sites flanking pUC18 polylinker | (Herrero *et al.*, 1990) |
| pUC18Not_ΔcheR | Ap^R^; 1.5-kb PCR product containing a 678 bp in frame deletion of *cheR* of *P. atrosepticum* inserted into the EcoRI/PstI sites of pUC18Not | This study |
| p34S-Km3 | Km^R^, Ap^R^; Km3 antibiotic cassette | (Dennis and Zylstra, 1998) |
| pUC18Not_ΔcheR_km3 | Ap^R^, Km^R^; 0.95-kb BamHI fragment containing Km3 cassette of p34S-Km3 was inserted into BamHI site of *cheR* in pUC18Not_ΔcheR | This study |
| pUC18Not-pacC-NWTTF | Ap^R^; 1-kb PCR product with NWETF pentapeptide of PacC mutated to NWTTF and inserted into the EcoRI/PstI sites of pUC18Not | This study |
| pUC18Not-pacC-GWTTF | Ap^R^; 1-kb PCR product with NWETF pentapeptide of PacC mutated to GWTTF and inserted into the EcoRI/PstI sites of pUC18Not | This study |
| pKNG101 | Sm^R^; *oriR6K mob sacBR* | (Kaniga *et al.*, 1991) |
| pKNG101_ΔcheR | Sm^R^, Km^R^; 2.5-kb NotI fragment of pUC18Not_ΔcheR_Km3 was cloned at the same site in pKNG101 | This study |
| pKNG-pacC-NWTTF | Sm^R^; 1.1-kb NotI fragment of pUC18Not-pacC-NWTTF cloned into the EcoRI/PstI sites of pKNG101 | This study |
| pKNG-pacC-GWTTF | Sm^R^; 1.1-kb NotI fragment of pUC18Not-pacC-GWTTF cloned into the EcoRI/PstI sites of pKNG101 | This study |

^a^Ap, ampicillin; Em, erythromycin; Km, kanamycin; Sm, streptomycin.

**Table S4) Oligonucleotides used in this study.**

| **Name** | **Sequence (5’-3’)** | **Purpose** |
| --- | --- | --- |
| pET28_CheR_Pec___f | taatGCTAGCATGAGCAAGATAAGAGTATTATGCGTTG | Construction of pET28b-CheR |
| pET28_CheR___Pec___r | taatGTCGACGCTCACCGAGAGCTGCTTAT |  |
| pUC18Not_cheR_up_f | taatgaattccgttcagaagaacgacacaggc | Construction of pUC18Not_ΔcheR |
| pUC18Not_cheR_up_r | taatggatccCCGGTCAACCATCTGCGTC |  |
| pUC18Not_cheR_down_f | taatggatcCCGTCGATTCGTCCCGATGC |  |
| pUC18Not_cheR_down_r | taatctgcagGCGCAATATAGGCATGTCCAGG |  |
| ECA1691-EcoRI_F | taatgaattcAGGGCAGGAGAGCAGGGC | Amplification of *pacC* |
| ECA1691-PstI_R | attactgcagcatatcaaaccgctctaatcttg |  |
| ECA1691-NWTTF_F | TGACAATTGGACCACCTTCTGAt | Construction of pUC18Not-pacC-NWTTF |
| ECA1691-NWTTF_R | aTCAGAAGGTGGTCCAATTGTCA |  |
| ECA1691-GWTTF_F | ATCAACTGACGGCTGGACCACCTTCTGAt | Construction of pUC18Not-pacC-GWTTF |
| ECA1691-GWTTF_R | aTCAGAAGGTGGTCCAGCCGTCAGTTGAT |  |
